# Protein structure shapes natural genetic variation and *de novo* adaptation

**DOI:** 10.64898/2025.12.19.695599

**Authors:** Christopher M. Jakobson, Alexandria Van Elgort, José Aguilar-Rodríguez, Daniel F. Jarosz

## Abstract

Protein structural context strongly influences the functional outcomes of mutations, but exploiting this intuition to interpret the consequences of natural variants has been limited by the paucity of structural information. Here, we show that simple molecular heuristics based on predicted protein structures–solvent accessibility and the local complexity of the protein fold–are strong predictors of protein-coding variation in genomes from yeast to plants to humans. Polymorphisms in structured and unstructured regions alike were subject to these constraints, revealing fine-grained selection across amino acid residues and secondary structures. The same analyses distinguished deleterious variants in datasets from existing deep mutational scans. Extending the metrics to *de novo* adaptation, we developed a massively parallel directed evolution platform that isolated hundreds of independent adapted lineages within days. Optimized experimental conditions revealed dozens of rapamycin-resistant alleles of the *FPR1*/FKBP gene in a single experiment–resistance mutations that matched predictions from our structural heuristics. Together, these data demonstrate the power of a biochemical perspective to contextualize the functional landscape of both existing and newly arising genetic variation.

## MAIN TEXT

Decades of experimental evidence from enzymatic biochemistry makes clear that molecular context shapes the function of mutations: key catalytic and other functional sites preferentially harbor variants of large effect, whereas other regions are much more labile^1^. Yet most population genetic studies to date do not account for protein structural context when assessing the functional impact of mutations^2^. Historically, this disconnect was primarily due to the paucity of structural information to contextualize natural variants: typically, at most a few hundred proteins in a proteome have been crystallized or otherwise structurally characterized (*e.g.*, by nuclear magnetic resonance)^3^. Indeed, although the role of protein structure in shaping the rate of protein evolution has long been appreciated^4^, the lack of protein structural information represented a major barrier to interpreting rare or *de novo* natural variants inaccessible to statistical population genetics.

Now, however, backbone protein structure predictions are now available for essentially all proteins based on residue covariation analysis (*e.g.* from the AlphaFold project^5^). This presents a newfound opportunity to combine rapidly emerging data from genomics and protein structure prediction with laboratory approaches to assess the function of many variants in parallel. The AlphaMissense project, for instance, has predicted the functional consequences of many human missense variants^6^. Yet predictions are unavailable, and cannot be generated using publicly available resources, for other organisms. We therefore sought to use molecular heuristics, readily calculated based on predicted structures, that would be broadly applicable across the vast array of proteomes analyzed by AlphaFold2. We have shown experimentally that structural parameters are useful to distinguish natural variants that have a functional impact on protein levels^7^, and others have used them to predict which sites in a protein evolve more rapidly^8–11^.

Here, we show that molecular context, as defined by simple metrics from protein structures, strongly shapes the spectrum of natural protein-coding variation. These metrics apply to ordered and unstructured regions alike and highlighted the importance of conserved molecular principles in shaping the naturally occurring genetic variation in the budding yeast *Saccharomyces cerevisiae*. Indeed, these structural properties are alone sufficient to predict allele frequencies in natural populations from yeast to the model plant *Arabidopsis thaliana* (thale cress) and even in humans. Yet even in the most comprehensive population-wide datasets, variation in any one gene is sparse, limiting the assessment of constraint within individual proteins. Moreover, cancers and pathogens often exploit adaptive solutions that are otherwise rarely observed. Therefore, we developed a massively parallel evolution assay to rapidly isolate dozens of independent adaptive mutations in the same gene product. This forward-looking experimental approach revealed that the same biochemical influences shape the spectrum of *de novo* adaptive variants as have shaped the variants currently present from fungi to humans.

## RESULTS

### Molecular signatures of structural constraint across extant variation in budding yeast

Genetics often relies on an understanding of what variation is missing in a population, whether to identify essential genes or catalytic residues. Building on this idea, and using intuition from protein engineering and structural and evolutionary biology, we hypothesized that two key factors shape the constraint on an amino acid residue within populations are (I) its proximity to the folded core of the protein (as represented by its accessible surface area, or A.S.A.) and (II) the number of other amino acid residues nearby, interactions with which would be disrupted by a mutation (as represented by the number of α-carbon atoms within 10Å, or packing density). To test our hypothesis, we first generated a database of all possible single-nucleotide polymorphisms (SNPs) in the reference proteome of the budding yeast *S. cerevisiae* [**Fig. 1A**] and calculated each structural parameter (A.S.A. and packing density) for all ∼ 2 × 10^7^ possible missense SNPs [**Fig. 1B**]. To assess constraint, we then calculated these parameters for every natural missense variant (*n* ∼ 5 × 10^5^) observed in the 1,011 Yeast Genomes collection, a panel of highly diverse budding yeast isolates from around the world^12^ [**Fig. 1C**]. Any systematic differences between the actual and simulated variant distributions across these metrics indicates evolutionary constraint, as variants absent from natural populations have likely been subject to purifying selection.

**Figure 1.**
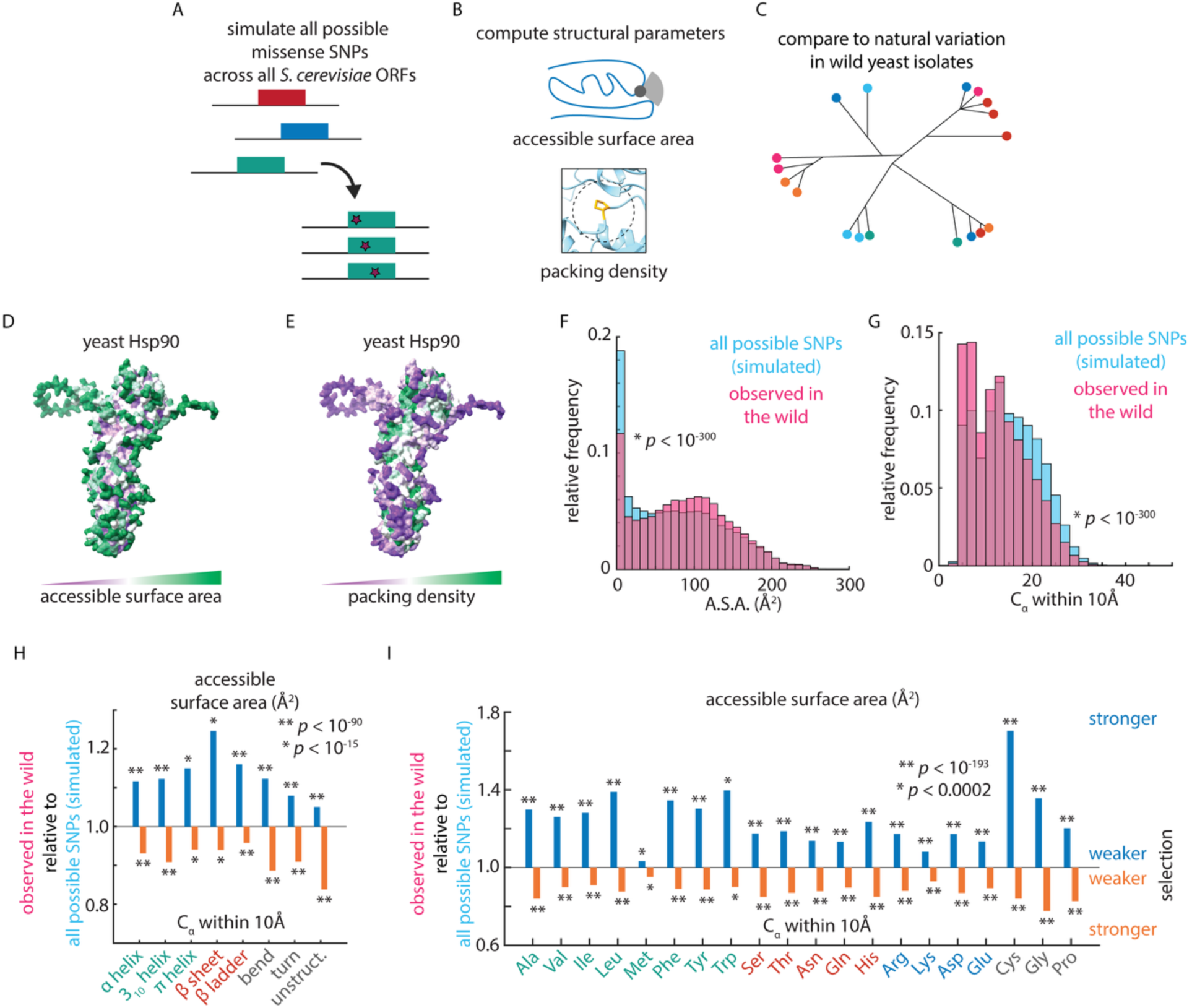
Structural constraint on natural genetic variation. (A) Simulation of all possible missense SNPs in the *Saccharomyces cerevisiae* proteome. (B) Computation of local protein structural parameters: solvent-accessible surface area (top) and local packing density (α-carbon atoms within 10Å). (C) Analysis of wild *S. cerevisiae* genome sequences. (D) Structure of *S. cerevisiae* Hsp90 (*HSP82*) with residues colored by accessible surface area, as shown. (E) As in (D), for packing density. (F) Relative frequency of accessible surface area for all possible missense SNPs in the *S. cerevisiae* proteome (blue) and missense SNPs observed in wild *S. cerevisiae* isolates (pink). *p* value by K.-S. test. (G) As in (F), for packing density. (H) Relative accessible surface area (blue) and relative packing density (orange) of missense SNPs observed in wild *S. cerevisiae* isolates as compared to all possible missense SNPs in the *S. cerevisiae* proteome, subset by local protein secondary structure, as indicated. *p* values by two-tailed *t* test. (I) As in (H), subset by reference amino acid identity as indicated.

As expected, these parameters varied over a wide range [**Fig. 1DE; Fig. S1AB**] and depended on both the reference amino acid and the local secondary structure in the wild-type fold [**Fig. S1CDEF**], which have their own typical biochemical characteristics. Indeed, naturally occurring missense variants were substantially more exposed to solvent (*p* < 10^−300^) [**Fig. 1F**] and resided near fewer neighboring amino acid residues (*p* < 10^−300^) [**Fig. 1G**] than all missense SNPs that could possibly have occurred. Fully buried residues (A.S.A = 0) were particularly rarely mutated amongst natural variants, whereas positions with low packing density (fewer than 10 close neighbors) harbored a clear excess of natural polymorphisms.

These enrichments were striking across all types of secondary structures and all wild-type amino acids, including those typically thought to be less subject to such constraint. Beta sheets and beta ladders, for instance, were highly constrained with respect to solvent accessibility (naturally variable residues were 15 - 25% more exposed to solvent; *p* < 10^−28^); unstructured regions were much less so (∼ 5% more solvent-exposed; *p* < 10^−300^) [**Fig. 1H**] – likely because these secondary structures are more prone to participate in the folded core of proteins. Likewise, for the same reason, hydrophobic residues were much more constrained in terms of packing density than charged residues [**Fig. 1I**]. Conversely, unstructured regions exhibited strong selection on the basis of the number of nearby residues (naturally variable residues had ∼ 15% fewer neighbors; *p* < 10^−300^) [**Fig. 1H**], as did glycine residues (> 20% fewer neighbors; *p* < 10^−200^) [**Fig. 1I**]. This may reflect the need to preserve local conformational flexibility or avoid steric clashes that disrupt transient interactions or dynamic conformational ensembles, which are often critical for protein function. These data suggest that even flexible regions are nonetheless subject to biochemical constraint that can be detected using protein structural heuristics.

### Influence of local structure on variation within and between proteins

The concordance of our findings with the fine details of protein structure led us to ask whether we could identify patterns of structural selection within the residue and secondary structure categories we examined. Strong signatures of purifying selection were evident, for instance, for glycine and cysteine residues in unstructured regions [**Fig. 2A**]. The latter likely stems from the unique chemistry made possible by the thiol group of cysteine residues, but we were surprised to observe such striking constraint on glycine residues that we had expected to be more labile. We therefore examined the dihedral angles (conventionally annotated as the Φ and Ψ angles [**Fig. 2B**]) of all possible *versus* naturally mutated glycine residues. Indeed, natural variants at glycine residues in unstructured regions were strikingly conformationally distinct from all possible mutations to such residues (*p* < 10^−24^; *p* < 10^−68^), suggesting that certain poses of glycine were more permissive to variation [**Fig. 2CD**]. Together, these results indicate that the structural heuristics we propose are a powerful metric to identify signals of selection both within and across residues and secondary structural contexts.

**Figure 2.**
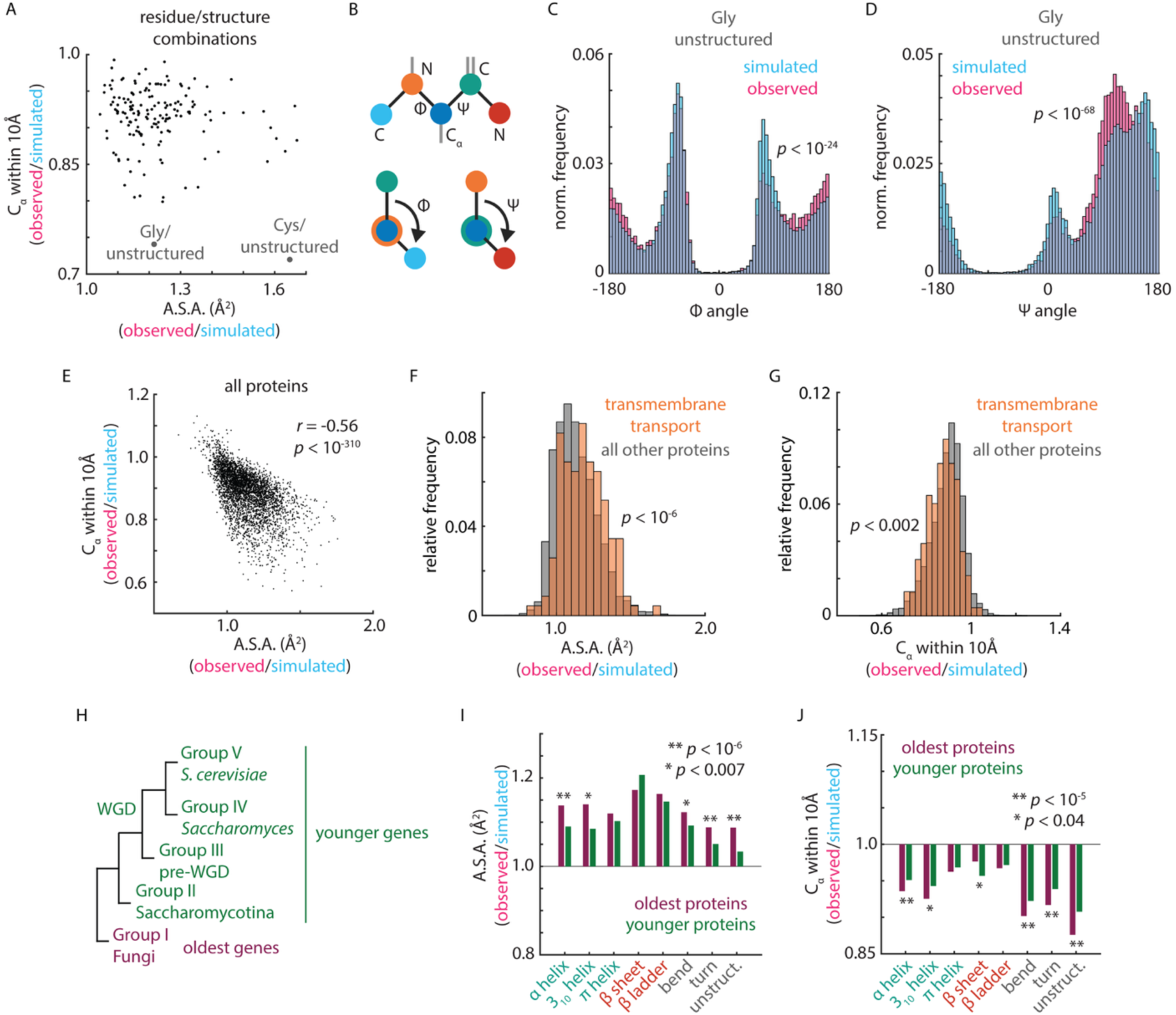
Structural and functional basis of constraint on coding variation. (A) Mean relative packing density (ordinate) and mean relative accessible surface area (abscissa) for missense SNPs observed in wild *S. cerevisiae* isolates relative to all possible missense SNPs for all combinations of local protein secondary structure and reference amino acid identity. Highlighted are glycine and cysteine reference residues in unstructured regions. (B) Schematic of Φ and Ψ angles in polypeptide chains. (C) Φ angle distribution for all possible missense SNPs at glycine residues in unstructured regions (blue) and missense SNPs observed at glycine residues in unstructured regions in wild *S. cerevisiae* isolates (pink). *p* value by K.-S. test. (D) As in (C), for Ψ angle. (E) Mean packing density (ordinate) and mean accessible surface area (abscissa) across all residues for each protein-coding gene in the *S. cerevisiae* proteome. *p* value from *t* distribution. (F)Mean relative accessible surface area for missense SNPs observed in wild *S. cerevisiae* isolates relative to all possible missense SNPs for transmembrane transporters (orange) and all other proteins (grey). (G) As in (F), for mean relative packing density. (H) Schematic of age stratification of *S. cerevisiae* genes by time of emergence. (I) Relative accessible surface area for missense SNPs observed in wild *S. cerevisiae* isolates as compared to all possible missense SNPs in the *S. cerevisiae* proteome, subset by secondary structure and by oldest (purple) and younger (green) genes, as indicated in (H). *p* values by *t* test. (J) As in (I), for relative packing density.

Recognizing that distinct functional classes of proteins (*e.g.*, enzymes *versus* structural components of the cytoskeleton *versus* membrane proteins) have very different amino acid and secondary structure content shaped by their activities and folds, we next calculated the average relative ASA and packing density across all residues for each protein in the *S. cerevisiae* proteome. As expected, given that proteins experience disparate strengths of selection, the relative solvent accessibility and packing density for naturally occurring variants were highly correlated on a gene-wise basis (*r* = −0.55, *p* < 10^−300^) [**Fig. 2E**]. These metrics were also well-correlated with the statistic *pN*^13^ (here, simply the number of naturally occurring missense variants relative to the number of possible missense SNPs), classically used to infer evolutionary constraint within populations^14^ (*r* = −0.47, *p* < 10^−212^; *r* = 0.34, *p* < 10^−106^) [**Fig. S2AB**]. The most constrained proteins were enriched in metabolism (*p* < 10^−16^), biosynthesis (*p* < 10^−11^), and transmembrane transport (*p* < 10^−10^) [**Fig. 2FG**], consistent with the central roles and highly conserved components of these processes^15^.

To assess the origins of the gene-wise variation in the strength of selection that we observed, we compared the apparent strength of selection on the most ancient proteins in *S. cerevisiae* (those shared across fungi, with a common ancestor dating to over 400 million years ago) to more novel proteins (found only in Saccharomycotina)^16^ [**Fig. 2H**], reasoning that ancient proteins might exhibit stronger evidence for selection because they are enriched for essentiality and have experienced a much longer duration of purifying selection. Indeed, ancient proteins showed stronger signals of selection across all three metrics [**Fig. 2IJ**; **Fig. S2CDEF**].

In addition to variation in overall structural context between proteins, many proteins contain well-folded catalytic or other functional domains alongside less-structured regions in the same polypeptide. To investigate the influence of this distinction, we used Superfam conserved domains^17^ as a proxy to identify folded, functional regions of proteins, and subdivided all residues in the proteome on this basis. Indeed, residues in Superfam domains had much lower background solvent accessibility (*p* < 10^−16^) and higher packing density (*p* < 10^−16^) [**Fig. S2G**] than other regions of proteins. Naturally occurring variants were more solvent accessible (*p* < 10^−16^, *p* < 10^−16^) and had lower packing density (*p* < 10^−16^, *p* < 10^−16^) both within and outside well-folded functional modules [**Fig. S2G**]; that is, the biochemical basis for selection we observed above was uniformly evident for both domain classes.

### Structural constraint across the continuum of natural variant frequencies

Having thus far compared all natural variants in a survey of budding yeast to all possible simulated missense SNPs, we reasoned that the same evolutionary forces might describe the frequency of variants within *S. cerevisiae* populations. In this case, systematic differences in solvent accessibility and packing density between rare and common natural variants would reveal ongoing purifying selection, as deleterious variants are expected to remain at low frequency or be eliminated from the population. To this end, we divided the naturally occurring missense variants into rare (allele frequency ≤ 1%) and common (allele frequency > 1%) variants. Indeed, common variants (*n* ∼ 91,000) were significantly more solvent-exposed (*p* < 10^−179^) [**Fig. 3A**] and occurred in less complex regions of the protein fold (*p* < 10^−160^) [**Fig. 3B**] than variants that were rare in nature (*n* ∼ 433,000). Moreover, the strength of selection was much more evident when we subdivided variants based on their wild-type secondary structure [**Fig. 3C**] and reference amino acid residue [**Fig. 3D**], both for solvent accessibility and the number of nearby residues. Indeed, as we observed above when comparing observed to all simulated variants, selection on solvent accessibility was most pronounced for beta ladders (*p* < 10^−14^); likewise, unstructured regions exhibited the strongest selection on packing density (*p* < 10^−76^) [**Fig. 3C**]. Consistent with their typically high solvent exposure and hydrophilicity, charged residues experienced the weakest strength of selection on solvent accessibility [**Fig. 3D**].

**Figure 3.**
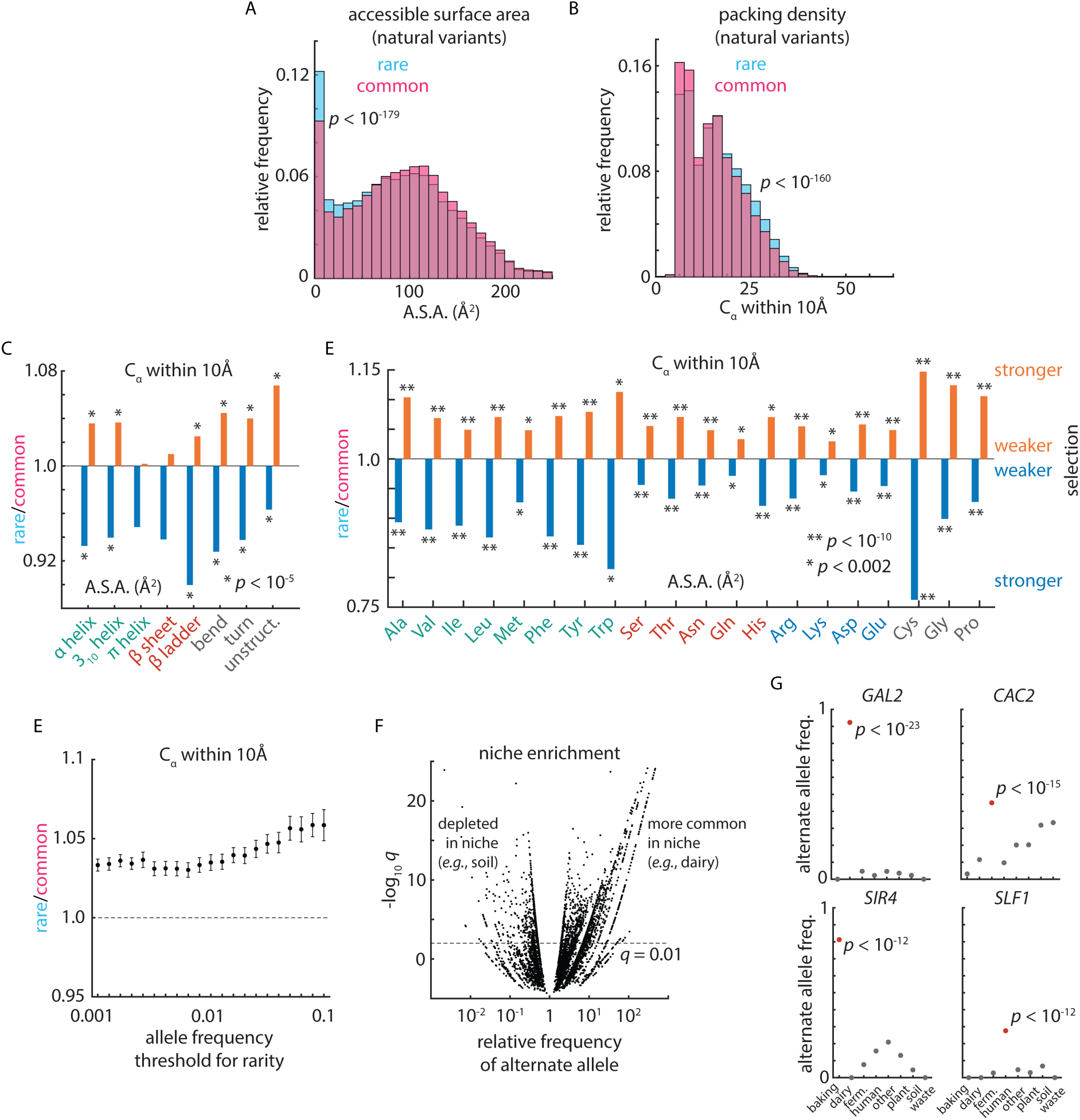
Signatures of selection across wild yeast. (A) Relative frequency of accessible surface area for rare (blue) and common (pink) missense variants observed in wild *S. cerevisiae* isolates. *p* value by K.-S. test. (B) As in (A), for packing density. (C) Relative accessible surface area (blue) and relative packing density (orange) of rare relative to common missense variants observed in wild *S. cerevisiae* isolates, subset by local protein secondary structure, as indicated. *p* values by *t* test. (D) As in (C), subset by reference amino acid identity as indicated. (E) Relative packing density of rare relative to common missense variants observed in wild *S. cerevisiae* isolates for a range of allele frequency thresholds for rarity, as indicated. (F) Volcano plot illustrating enrichments of natural alleles in ecological niches, showing the relative enrichment of the minor allele (abscissa) and the Bonferroni-corrected Fisher’s exact test *q* value (ordinate). (G)Example enrichments of natural alleles in niches as indicated.

Although variants are often categorized as rare based on the 1% allele frequency threshold used above, we wondered whether signatures of selection might be evident across a wider range of definitions. We therefore calculated the relative solvent accessibility and number of nearby residues for rare *versus* common natural variants across a range of minor allele frequencies from 0.1% to 10% [**Fig. 3E**; **Fig. S3A**]. Strikingly, the impress of selection was evident across this range of frequencies, particularly for packing density, suggesting that these molecular metrics reflect pervasive selection across rare and common variation.

We also noted that, despite these pronounced signatures of constraint, many common variants (allele frequency > 5%) were nonetheless biochemically perturbative. We therefore asked whether their prevalence could be explained by positive selection on their adaptive benefits in particular ecological niches. Indeed, classifying wild yeast isolates into eight broad niche classes (baking, dairy, fermentation, human-associated, plant, soil, waste, and all other) revealed that many common variants bore striking enrichments in one niche as compared to the others (> 6,000 of ∼ 21,000 common alleles with allele frequency > 5% had significant niche enrichment; *q* < 0.01) [**Fig. 3F**]. These included variants in the galactose permease *GAL2* (*q* < 10^−24^; associated with dairy isolates), the chromatin assembly factor *CAC2* (*q* < 10^−16^; fermentation strains), the silencing regulator *SIR4* (*q* < 10^−13^; baking yeasts), and the RNA-binding protein *SLF1* (*q* < 10^−13^; human-associated isolates) [**Fig. 3G**]. Indeed, a prior study identified balancing selection on *GAL2* linked to galactose-rich environments^18^. Thus, common variation in *S. cerevisiae* is replete with signatures of adaptive evolution and biochemical constraint—signatures that are clearer in light of both the ecological and protein structural context of natural variants.

### Molecular heuristics predict natural variation and fitness effects across Eukarya

Although a powerful model organism for laboratory and ecological studies, the effective population size in budding yeast (conservatively, *N_e_* ∼ 10^7^ or larger), and thus the strength of selection, is likely much greater than in plants or animals^19^. We therefore sought to test our heuristics in other species spanning a range of effective population sizes: the thale cress *Arabidopsis thaliana* (*N_e_* ∼ 10^5^)^20^ and *Homo sapiens* (*N_e_* ∼ 10^4^ or smaller)^21^. First, in *A. thaliana*, we made use of the 1,001 Genomes catalogue^22^ and mapped the missense variants from these accessions (*n* ∼ 1.1 × 10^6^) onto the AlphaFold2 backbone structure predictions for the *A. thaliana* proteome. The variants identified in this collection span a comparable range of allele frequencies to those we examined in *S. cerevisiae* [**Fig. S4A**]. Comparing rare (allele frequency ≤ 0.1%) to common (> 0.1%) revealed that rare variants were indeed more buried (*p* < 10^−183^) [**Fig. 4A**] and had more nearby residues (*p* < 10^−104^) [**Fig. 4B**] than common variants. As in budding yeast, this was true across protein secondary structures [**Fig. 4C**]. Thus, the patterns we identified above were not due solely to the exceptionally high selective pressures on microbes.

**Figure 4.**
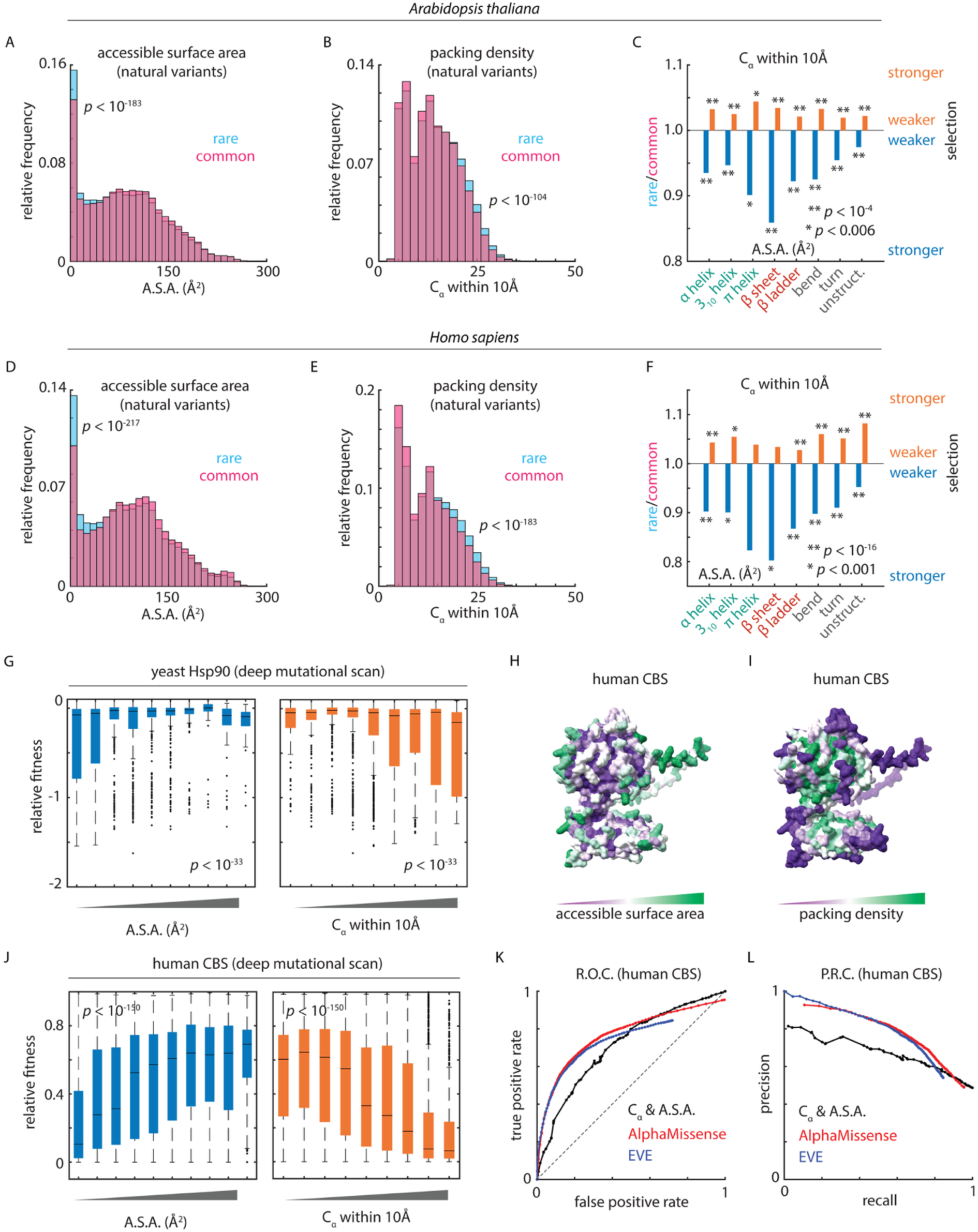
Molecular heuristics predict coding variant frequencies and fitness effects across Eukarya. (A) Relative frequency of accessible surface area for rare (blue) and common (pink) missense variants observed in wild *A. thaliana* accessions. *p* value by K.-S. test. (B) As in (A), for packing density. (C) Relative accessible surface area (blue) and relative packing density (orange) of rare relative to common missense variants observed in wild *A. thaliana* accessions subset by local protein secondary structure, as indicated. *p* values by *t* test. (D-F) As in (A-C), for natural human genetic variation in gnomAD. (G) Relative fitness (ordinate) of synthetic missense variants of *S. cerevisiae* Hsp90 (*HSP82*) as a function of accessible surface area (left) and packing density (right). *p* values by *t* statistic of Spearman correlation. (H) Modeled structure of *H. sapiens* cystathionine beta-synthase (CBS) with residues colored by accessible surface area, as shown. (I) As in (H), for packing density. (J) Relative fitness (ordinate) of synthetic missense variants of *H. sapiens* CBS as a function of accessible surface area (left) and packing density (right). *p* values by *t* statistic of Spearman correlation. (K) Receiver operating characteristic curve for classification of deleterious CBS variants based on structural parameters (black), AlphaMissense (red), and EVE (blue). (L) Precision-recall curve for classification as in (K).

Next, we exploited the gnomAD database of human genetic variation^23^ and charted the missense variants (*n* ∼ 4.4 × 10^6^) from this collection onto the predicted structures of the human proteome. This resource contains relatively more rare variants than the yeast and plant populations we examined above but nonetheless harbors a wide range of more common variation [**Fig. S4A**]. Remarkably, considering the evolutionary distances and disparities in effective population size, we obtained very similar results in human as in yeast and *A. thaliana* when considering rare (allele frequency ≤ 0.1%) *versus* common (> 0.1%) missense variants: rare variants were once again less solvent-exposed (*p* < 10^−217^) [**Fig. 4D**] and arose in more complex regions of the protein fold (*p* < 10^−183^) [**Fig. 4E**]. This was true across well-ordered secondary structures, such as alpha helices and beta sheets, as well as in unstructured regions, which make up a large fraction of the proteome in all three organisms [**Fig. 4F**; **Fig. S4B**].

Lastly, although they are powerful resources to assess the action of selection in the wild, catalogues of natural genetic variants are typically sparse for any single gene (median 69, 31, and 184 missense SNPs per gene in the *S. cerevisiae*, *A. thaliana*, and *H. sapiens* databases we analyzed, respectively) [**Fig. S4C**]. For this reason, studies quantifying constraint typically aggregate variants from throughout a genomic window^23^, precluding detailed investigation of where coding variants are permissible within an individual protein. We therefore turned to comprehensive deep mutational scans of the protein chaperone Hsp90 from *S. cerevisiae*^24^ and the essential metabolic enzyme cystathionine beta-synthase (CBS) from humans^25^. *In vitro e*xperimental data for CBS are highly predictive of pathogenicity and correlated with a range of patient outcomes, providing a well-validated and comprehensive test case. After calculating the relevant structural heuristics for each residue, we once again found that both solvent accessibility and packing density were strikingly predictive of highly deleterious variants: buried residues and those in crowded regions were much more likely to have large fitness consequences when mutated [**Fig. 4GHIJ**]. Classification of CBS mutant fitness based on these simple metrics performed surprisingly well compared to complex prediction algorithms, achieving a true positive rate of 53% at 25% false positive rate, while EVE^26^ and AlphaMissense^6^ achieved true positive rates of 68% and 69%, respectively, at this threshold [**Fig. 4K**]. Discriminating moderately deleterious variants remained challenging even for these cutting-edge models: all three approaches had similar precision at 75% recall (A.S.A. and packing density: 61%; EVE: 67%; AlphaMissense: 70%) [**Fig. 4L**]. Thus, these molecular heuristics capture important aspects of selection on coding variation throughout Eukarya and likely represent a valuable tool for understanding the consequences of natural variants across diverse non-model organisms.

### Highly parallelized mutation-limited laboratory evolution

A powerful complement to deep mutational scanning, which typically targets a single gene, is laboratory directed evolution, which can identify mutations throughout the genome that converge on coherent pathways and functions^27^ and provides an avenue to identify functional variation *en masse*. Yet, as classically performed, this approach also surveys only a limited spectrum of the distribution of adaptive mutations, largely due to clonal interference within a limited number of large populations^28–30^. Moreover, experiments in chemostats are known to be susceptible to selection for undesired phenotypes, such as aggregation^31^. To circumvent these limitations, we employed a massively parallel directed evolution paradigm based on thousands of independent, mutation-limited populations of *S. cerevisiae* [**Fig. 5A**]. We reasoned that this platform, under appropriate conditions, would allow us to rapidly identify many adaptive mutations in a single gene product or pathway.

**Figure 5.**
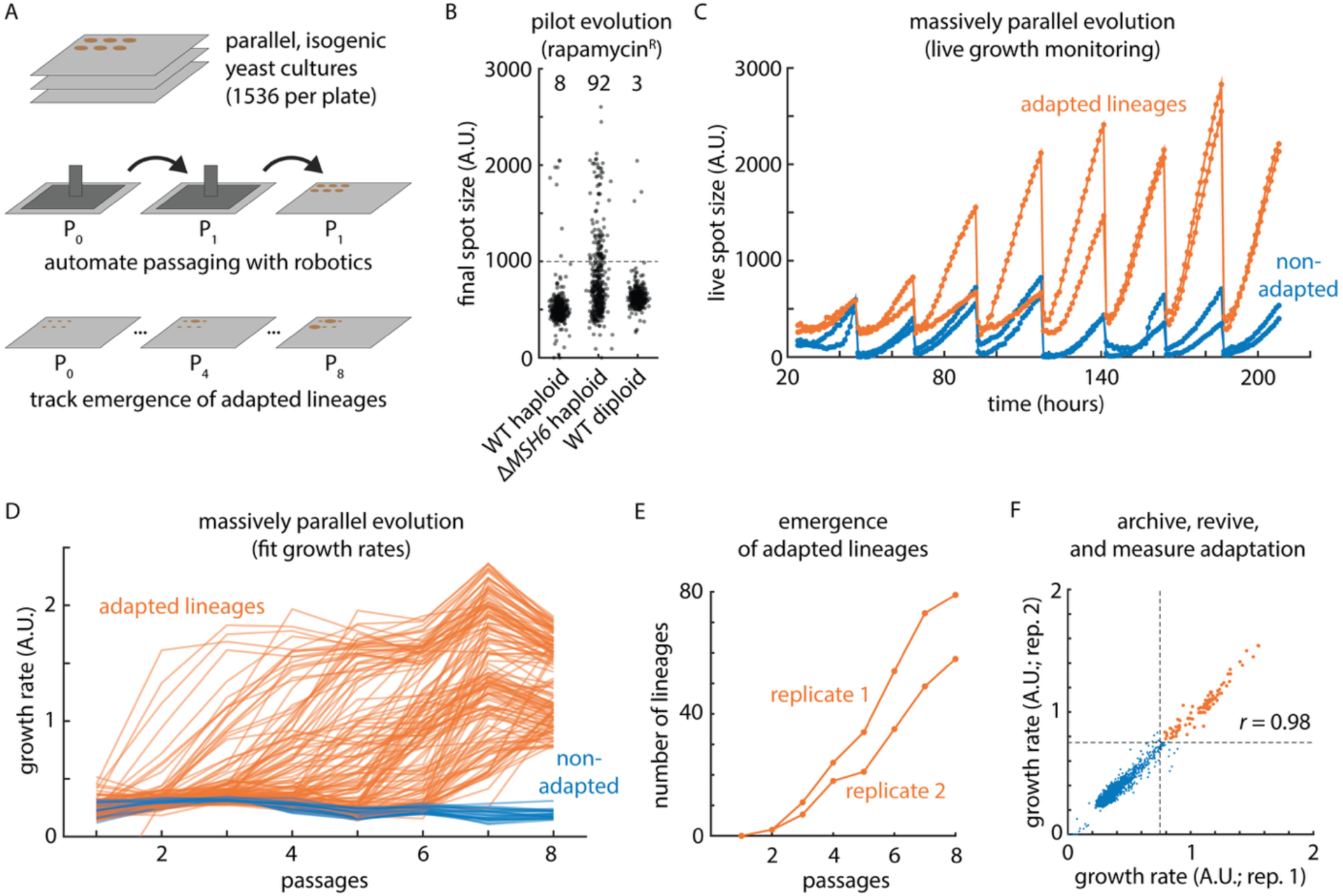
A massively parallel platform for experimental evolution. (A) Schematic of massively parallel microbial directed evolution platform. (B) Final spot sizes in pilot rapamycin-resistance evolution campaign for BY4741 genotypes and ploidies as indicated. Noted are the number of strongly adapted lineages. (C) Exemplary continuous spot size measurements for adapted (orange) and non-adapted (blue) lineages. (D) Per-passage growth rate estimates for all evolving lineages across primary rapamycin-resistance evolution campaign. Lineages that adapted by passage 8 are colored orange. (E) Rate of emergence of adapted lineages in independent replicate plates. (F) Replicability of estimated growth rates in rapamycin after archiving and revival of directed evolution lineages.

We chose to evolve mutants resistant to the natural product rapamycin^32^ because its mode of action – inhibition of the key TOR signaling complex – is deeply characterized^33^ and because a pioneering 1991 study isolated nine resistant clones due to mutations in *FPR1* (FKBP; the target, with Tor, of rapamycin)^34^. We hypothesized that adaptation to this environment would be less genetically complex than evolution in, for instance, rich medium – a condition in which adaptive variants can arise in a relatively large number of genes^27,35^. As a proof-of-concept, we passaged 384 isogenic haploid laboratory yeast populations every 24 hours on rich YPD solid medium with 10 nM rapamycin for a total of 8 days. At the end of this regimen, we identified 8 strongly adapted lineages (final spot size > 1000 A.U.; ∼ 2% of the starting populations) [**Fig. 5B**]. To test whether the evolved populations were indeed mutation-limited in their capacity for adaptation, we included an equal number of lineages lacking the mismatch repair factor *MSH6*. Cells lacking Msh6 exhibit up to 50-fold higher mutation rates^36^ and indeed, as expected in the mutation-limited regime, this arm of the evolution campaign yielded 92 adapted lineages [**Fig. 5B**]. Finally, the same number of evolving wild-type diploid lineages yielded only 3 adapted lineages, consistent with the expectation that many loss-of-function resistance mutations in drug targets are recessive^37^ [**Fig. 5B**].

### Massively parallel directed evolution identifies the spectrum of rapamycin-resistant variants

Building on this proof of principle, we conducted a large directed evolution campaign under the same conditions (YPD + 10 nM rapamycin), evolving 1536 initially isogenic populations in each of two biological duplicate arms (3072 total lineages) [**Fig. 5C**]. Continuous growth monitoring by optical scanning throughout the experiment allowed us to readily identify a total of 137 strongly adapted lineages (58 and 79 from each respective arm) [**Fig. 5D**], which emerged at a similar rate throughout the experiment (∼ 17 lineages, or 0.5%, per passage) [**Fig. 5E**]. To confirm the accuracy of the *in situ* growth rate measurements used to track adaptation, we archived the final passaged populations after we ceased the directed evolution campaign and measured their growth rates in duplicate. These replicate measurements agreed well (*r* = 0.98) [**Fig. 5F**]. As a control for the possibility of spontaneous rapamycin resistance emerging in the absence of selection, we also passaged 1536 strains in medium without drug for the duration of the experiment; these lineages never exhibited resistance [**Fig. S5A**]. We also confirmed that the adapted lineages did not generally exhibit higher growth rates in media without drug (*p* = 0.06) [**Fig. S5B**].

To test our hypothesis that our approach would identify adaptive (*i.e.* loss-of-function) mutations in *FPR1*, we collected eight strongly adapted clones [**Fig. 6A**] and subjected them to whole-genome sequencing. As controls, we also sequenced the genomes of eight lineages that had been passaged for the same amount of time in media without drug; these isolates bore no fixed mutations relative to their ancestor. Four of the eight rapamycin-resistant lineages, however, bore mutations in *FPR1* [**Fig. 6B**; **Table 1**; **Table S1**]. We also identified a coding SNP in *P*leiotropic *D*rug *R*esistance 1 (mutations in *PDR1* are known to impact rapamycin resistance^38^). Intriguingly, we did not identify any fixed SNPs in three of the adapted lineages. This could, in some cases, be due to late-arising mutations that did not fix; below we discuss possible non-genetic mechanisms of resistance that might also be at play in these lineages. These data indicated that our massively parallel laboratory evolution platform was a powerful tool to rapidly identify mechanistically coherent resistance alleles.

**Figure 6.**
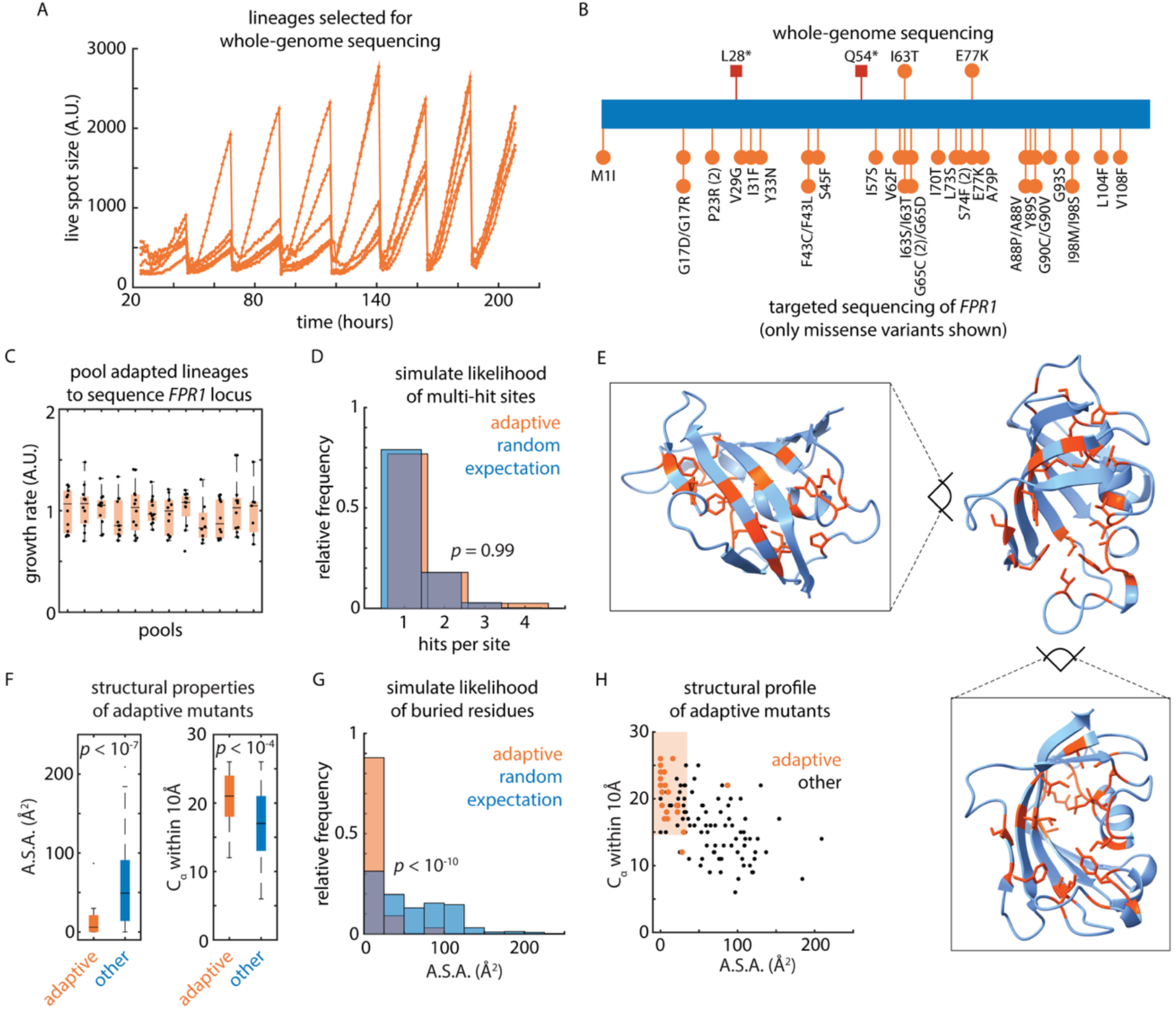
Directed evolution charts the biochemical solutions to drug challenge. (A) Growth trajectories of adapted lineages selected for whole-genome sequencing. (B) *FPR1* genotypes of populations subjected to whole-genome sequencing (top) and populations subjected to targeted *FPR1* amplicon sequencing (bottom). Only missense variants from amplicon sequencing are shown. (C) Growth rates of pools of adapted populations selected for targeted *FPR1* amplicon sequencing. (D) Relative frequency of multiple-hit sites in Fpr1 amongst adapted lineages (orange) and under simulated random mutations (blue). *p* value by K.-S. test. (E) Illustration of positions of missense variants identified in adapted lineages (orange) in the context of the predicted backbone structure of Fpr1 (blue). (F) Accessible surface area (left) and packing density (right) for missense variants identified in adapted lineages (orange) and all other residues in Fpr1 (blue). *p* values by *t* test. (G) Relative frequency of accessible surface area for missense variants identified in adapted lineages (orange) and under simulated random mutations (blue). *p* value by K.-S. test. (H) Profile of accessible surface area (abscissa) and packing density (ordinate) for missense variants identified in adapted lineages (orange) as compared to all other residues in Fpr1. Highlighted is the region of A.S.A. < 30 Å^2^ and more than 15 close neighbors.

**Table 1.**
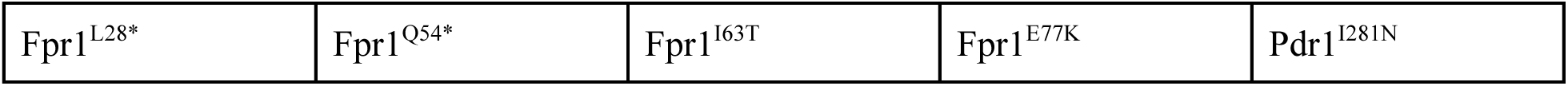
Fixed coding SNPs identified in whole-genome sequencing of rapamycin-resistant clones. Three other lineages that we sequenced had no fixed SNPs.

|  |  |  |  |  |
| --- | --- | --- | --- | --- |
| Fpr1<sup>L28*</sup> | Fpr1<sup>Q54*</sup> | Fpr1<sup>I63T</sup> | Fpr1<sup>E77K</sup> | Pdr1<sup>I281N</sup> |

### Protein structural context shapes de novo adaptation

To conduct a census of possible adaptive solutions in the experiment, we gathered 135 independent rapamycin-adapted lineages (12 pools of up to 12 different adapted lineages, selected in descending order of final adapted spot size) and subjected them to targeted deep sequencing of the *FPR1* locus [**Fig. 6C**]. We identified a total of 33 coding variants, as well as 29 indels and 18 premature termination codons and other presumed loss-of-function variants [**Fig. 6B**; **Table 2**; **Table S2**]. All told, at least 65.1% of sequenced adapted lineages bore an *FPR1* mutation, and these encompassed all four *FPR1* alleles identified by whole-genome sequencing. Moreover, we recovered three *FPR1* missense variants, two lineages abolishing the start codon of *FPR1*, and two lineages with premature stop codons that were all identified in the original study of spontaneous rapamycin-resistant mutants^34^; four of these were recovered in more than one independent isolate in our hands [**Table 2**].

**Table 2.** Missense mutations identified in pooled, targeted sequencing of the *FPR1* locus. Mutations also identified in whole-genome sequencing [**Table 1**] in **bold**. Mutations isolated in the 1991 identification of rapamycin resistant *FPR1* mutants^34^ <u>underlined</u>. See **Table S2** for other presumed loss-of-function mutants identified in targeted *FPR1* sequencing.

|  |  |  |  |  |  |  |
| --- | --- | --- | --- | --- | --- | --- |
| G17D | G17R | <u>P23R</u> (x2) | V29G | I31F | Y33N | F43C |
| F43L | S45F | I57S | V62F | I63S | **I63T** | <u>G65C</u> (x2) |
| <u>G65D</u> | I70T | L73S | S74F (x2) | **E77K** | A79P | A88P |
| A88V | Y89S | G90C | G90V | G93S | I98M | I98S |
| L104F | V108F |  |  |  |  |  |

At first blush, the resistance mutations we recovered within *FPR1* appeared nearly random: they were distributed throughout the coding sequence [**Fig. 6B**] and the rate of multiple mutations fixing at the same site was no higher than expected by chance for this sample size [**Fig. 6D**]. Striking commonalities between the mutations emerged, however, when we examined them in the context of the three-dimensional structure of Fpr1: practically all of the mutations were buried in the core of the folded domain, with many lying near the rapamycin-binding pocket [**Fig. 6E**]. Indeed, the Fpr1 variants we isolated were far less solvent-exposed (*p* < 10^−7^) and had much higher packing density (*p* < 10^−4^), than all possible missense SNPs in this protein [**Fig. 6F**] – that is, they resided precisely where loss-of-function variants would be predicted to arise by our population genetic analyses.

In contrast to the coarse-grained distribution of variants throughout the linear open reading frame, this pattern was highly unlikely to have occurred by chance amongst the lineages we collected (*p* < 10^−10^; *n* = 1,000 simulations) [**Fig. 6G**]. Perhaps most remarkably, we recovered lineages representing mutations to 21 out of 42 residues with this biochemical profile within the Fpr1 protein (A.S.A. < 30 Å^2^ and more than 15 close neighbors; 50%) [**Fig. 6H**]. In several cases, the same residue was mutated more than once, sometimes to more than one destination amino acid. Thus, massively parallel directed evolution for organismal phenotypes can achieve excellent coverage of resistance mutations in a single gene product, revealing that the same biochemical principles shape both these *de novo* adaptive mutations and the natural genetic variation segregating in a species as a whole.

## DISCUSSION

Understanding the consequences of naturally occurring variants remains a complex and challenging problem. Recent progress in classifying, for instance, deleterious variants in humans^6^ depends not only on proprietary algorithms, but also on extensive existing data on the effects of individual variants for training and refinement. Here, we demonstrate that leveraging ubiquitous protein structure predictions^5^, combined with simple structural calculations of solvent accessibility and packing density, offers a lens to understand selection on natural variation across species, predict the results of deep mutational scans, and appreciate the origins of adaptive variants that arise *de novo* under directed evolution. These analyses revealed common biophysical principles governing different secondary structures and reference amino acid residues, with strong signatures of selection evident even for conformationally flexible residues and unstructured regions.

Despite radically different effective population sizes, and likely disparate strengths of selection, the same trends in solvent accessibility and packing density held across catalogues of natural genetic variation in fungi, plants, and humans. In turn, the same approach will likely prove useful in interpreting the functional consequences of variants in microbes with few sequenced isolates and for metagenome-derived protein sequences that lack a reference genome. Notably, biophysical metrics distinguished not only standing genetic variation writ large from all mutations that could have occurred, but also rare from common variants in diverse datasets. This indicates that these heuristics describe fine details of selection, not simply coarse purifying selection against highly deleterious variants. Moreover, many highly perturbative common variants in *S. cerevisiae* bore striking signals of enrichment in a particular ecological niche, indicating the possibility of directional selection for adaptive variants under those conditions.

A key limitation of the population genetic datasets we initially analyzed is the relative sparsity of variants in any one gene product. To confirm the predictive power of our approach, and as a complement to these retrospective analyses, we also classified saturating mutations in yeast Hsp90 and human cystathionine beta-synthase (CBS) based on their fitness consequences. As expected, deleterious variants tended to be highly biophysically perturbative. Indeed, in the case of CBS, comparing our predictions to those of two elaborate classifiers, EVE and AlphaMissense, revealed only a modest deficit at modest rates of false discovery and comparable performance at high recall. This was without training or regression on mutagenesis data or clinical variant effect databases, nor even consideration of the reference or destination residue of each variant. Thus, much of the effect of a missense variant is encoded, and can be interpreted, merely on the basis of its context in a predicted protein structure – which we emphasize is now available for nearly all proteins in UniProt, in contrast to predictions of variant effects.

As a final test of this approach to missense variant interpretation, we devised a new massively parallel directed evolution paradigm to gather large numbers of adapted yeast lineages in high throughput. Exploiting this new technology allowed us to isolate dozens of *bona fide* drug-resistance variants in a single gene product (Fpr1) in only a week, and subjecting these mutations to structural analysis confirmed that solvent accessibility and packing density delineated the spectrum of possible adaptive variants – fully half of which we identified *in vivo*. We anticipate applying this platform to a variety of problems in evolutionary biology that would profit from a broad, unbiased survey of the mechanisms of adaptation^39^, including, for instance, the role of prions^40^ in mutagenesis and the emergence of new phenotypes^41^.

Our results illustrate the broad and straightforward applicability of protein structure to understanding existing and newly arising genetic variation, both in the laboratory and in the wild. In turn, this highlights a route toward biochemical variant interpretation across the vast majority of species and genera that lack genetic tools, collections of natural isolates, and assays of variant effect.

## METHODS

### Reference genomes and proteomes

The reference genome and transcriptome for *Saccharomyces cerevisiae* were S288C R64-1. The AlphaFold2 predictions for S*accharomyces cerevisiae*, *Arabidopsis thaliana*, and *Homo sapiens* were downloaded from https://www.alphafold.ebi.ac.uk/download. The *Arabidopsis thaliana* 1,001 Genomes genotype data was downloaded from https://1001genomes.org/data-center.html. GnomAD v2.1.1 genotype data (GRCh37/hg19 reference) were downloaded from http://www.gnomad-sg.org/downloads.

### Structural analysis of predicted protein structures

AlphaFold predicted backbone structures were analyzed with DSSP to derive secondary structure classifications, phi and psi angles, and solvent-accessible surface area for each residue. Custom code was used to calculate the number of alpha-carbons with 10Å of each alpha-carbon in each structure based on the PDB file coordinate. To simulate all possible SNPs in the S288C reference genome, we generated tables of all possible single-nucleotide variants, classified these as transitions or transversions, translated the mutated transcripts, and classified the mutations as missense or synonymous. Structural parameters for the mutated residues were tabulated with reference to the structural analyses of each protein described above.

### Multiplexed assays of variant effect data

MAVE^42^ data for CBS^25^ and Hsp90^24^ were downloaded from https://www.mavedb.org/score-sets/urn:mavedb:00000005-a-4 and https://www.mavedb.org/score-sets/urn:mavedb:00000074-a-1, respectively. AlphaMissense and EVE predictions for CBS were downloaded from https://www.alphafold.ebi.ac.uk/entry/P35520 and https://evemodel.org/download/protein/CBS_HUMAN, respectively.

### Yeast strains and growth conditions

Unless otherwise noted, yeast were propagated at 30℃ in yeast extract, peptone, and dextrose medium (10 g/L yeast extract; 20 g/L peptone; 20 g/L dextrose) with 20 g/L bacteriological agar added for solid media. Frozen stocks were stored with the addition of 15% glycerol and revived by pinning or streaking to solid medium.

### Massively parallel directed evolution

Yeast strains were revived from frozen stocks and cultivated in 384 or 1536 independent liquid precultures before robotic pinning using a Singer ROTOR to solid YPD medium without drug. These source plates were replicated to experimental conditions using robotic pinning and grown at 30℃ *in situ* on Epson V800 document scanners. Custom code was used to automate the scanners, image processing, and spot size quantification using the *gitter* R package^43^ (https://github.com/omarwagih/gitter/tree/master/R). Spots were passaged at 24-hour intervals and replaced on the scanning apparatus. At the conclusion of the directed evolution campaign, if desired, the fittest spots were identified and consolidated into 384-well plates for storage using a Singer PIXL colony picking robot.

### Whole-genome sequencing and genome analysis

Genomic DNA was prepared by growing cells to saturation in 1mL YPD, harvesting by centrifugation, and extracting DNA using a Norgen Fungi DNA Isolation kit #26200 according to the manufacturer’s directions. Library preparation used a NEB Next Ultra II FS library prep kit according to the manufacturer’s directions, with the exception that reactions were scaled down to yield approximately 6 μL of 5 ng/μL fragmented DNA per sample. Samples were barcoded using NEBNext Multiplex Oligos for Illumina 96 Unique Dual Index primers and submitted for sequencing with BGI on a DNBSEQ-G400. Trimmed reads were pseudo-aligned with *kallisto* and variants were called with respect to R64-1 using *bcftools*. Subtelomeric variants (within 30,000 bp of the chromosome end) were masked and variants were filtered based on QUAL > 150 and depth > 20 counts using custom code. Variants were functionally annotated using *sift*^44^. Sequencing reads are deposited at the Sequence Read Archive as BioProject PRJNA1289168.

### Pooled targeted sequencing

To genotype the *FPR1* locus, pools of adapted lineages were subjected to genomic DNA extraction as described above. This material was used as a PCR template to generate pooled amplicons from the *FPR1* locus. This mixed pool of products was deep-sequenced by Primordium sequencing (now Plasmidsaurus; https://plasmidsaurus.com/) and indels and SNPs collated using custom code.

### Data and code availability

All data required to reproduce the analyses and figures shown here are available at https://zenodo.org/records/17246894. WGS data is available at SRA (PRJNA1289168). Custom code to process and plot the data is available at https://github.com/cjakobson/pop-gen-structure. All strains are available upon request to Prof. Dan Jarosz.

## ACKNOWLEDGMENTS

We are grateful to all members of the Jarosz Lab for technical assistance and stimulating discussions.

## Funding

National Institutes of Health (DP2-GM119140, RF1-AG057334, R01-AG06341801, and R01-HG012366 to DFJ; F32-GM125162 to CMJ; T32-GM113854 to AV)

National Science Foundation (NSF-MCB116762 to DFJ; NSF GRFP to AV)

Searle Scholar Award (14-SSP-210 to DFJ)

Kimmel Scholar Award (SKF-15-154 to DFJ)

Vallee Scholar Award (to DFJ)

Discovery Innovation Award from Stanford University (to DFJ)

European Molecular Biology Organization (ALTF 724-2018 to JAR)

Swiss National Science Foundation (P2ZHP3_174735 to JAR)

DFJ is also a Science and Engineering Fellow of the David and Lucile Packard Foundation

Some of the computing for this project was performed on the Sherlock cluster. We would like to thank Stanford University and the Stanford Research Computing Center for providing computational resources and support that contributed to these research results.

## Author Contributions

Conceptualization: CMJ, AV, JAR, DFJ

Methodology: CMJ, AV

Investigation: CMJ, AV, JAR

Software, Formal Analysis, and Visualization: CMJ

Writing – Original Draft: CMJ

Writing – Review & Editing: CMJ, AV, JAR, DFJ

Supervision and Funding Acquisition: DFJ

