## Supplemental Information for "Protein structure shapes natural genetic variation and *de novo* adaptation"

Figures S1-S5

Tables S1 & S2

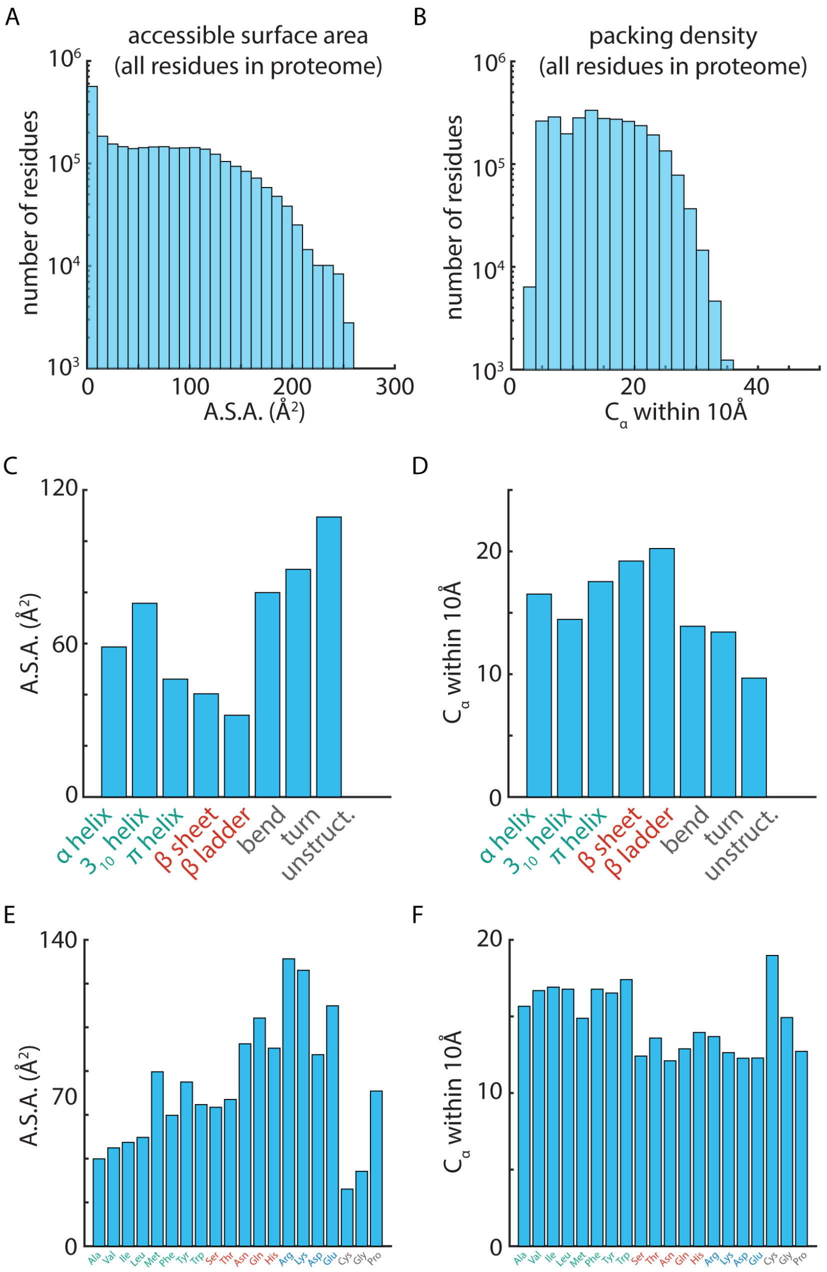


**Extended Data Figure S1. To accompany Figure 1.** (A) Frequency of accessible surface area for all residues in the *S. cerevisiae* proteome. (B) As in (A), for packing density. (C) Frequency of accessible surface area for all residues in the *S. cerevisiae* proteome, subset by local protein secondary structure, as indicated. (D) As in (C), for packing density. (E) As in (D), subset by reference amino acid identity as indicated. (F) As in (E), for packing density.

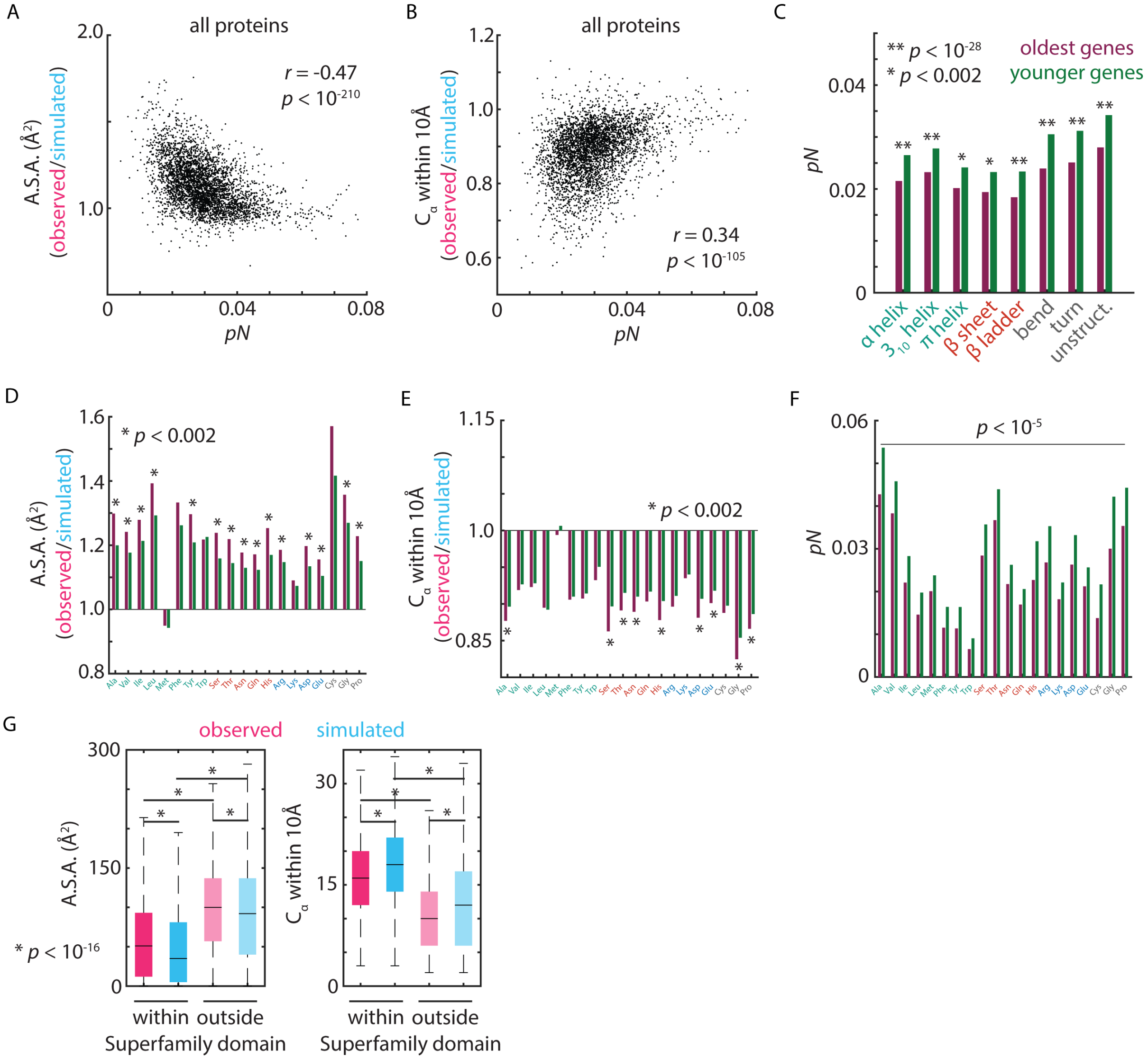


**Extended Data Figure S2. To accompany Figure 2.** (A) Mean accessible surface area (ordinate) and fraction of possible missense substitutions observed per-site *pN* (abscissa) across all residues for each protein-coding gene in the *S. cerevisiae* proteome. *p* value from *t* distribution. (B) As in (A), for mean packing density and *pN*. (C) Fraction of possible missense substitutions observed per-site *pN*, subset by secondary structure and by oldest (purple) and younger (green) genes, as indicated. *p* values by *t* test (D) Relative accessible surface area for missense SNPs observed in wild *S. cerevisiae* isolates as compared to all possible missense SNPs in the *S. cerevisiae* proteome, subset by reference amino acid identity and by oldest (purple) and younger (green) genes, as indicated. *p* values by *t* test (E) As in (D), for packing density. (F) As in (D), for *pN*. (G) Accessible surface area for all possible missense SNPs in the *S. cerevisiae* proteome (blue) and missense SNPs observed in wild *S. cerevisiae* isolates (pink), subset by positions within and outside SuperFamily domains, as indicated. *p* value by *t* test.

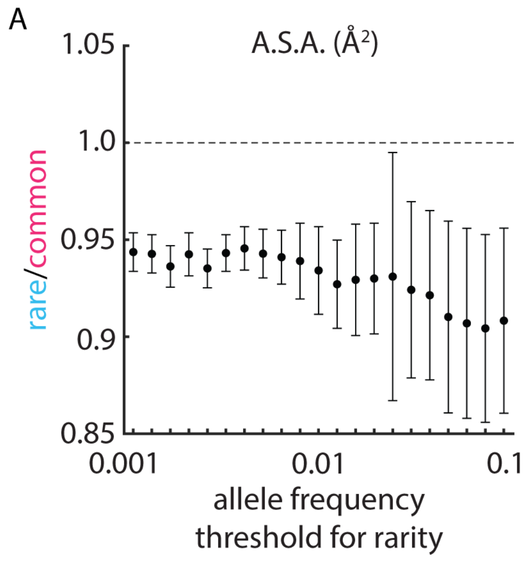


**Extended Data Figure S3. To accompany Figure 3.** (A) Relative accessible surface area of rare relative to common missense variants observed in wild *S. cerevisiae* isolates for a range of allele frequency thresholds for rarity, as indicated.

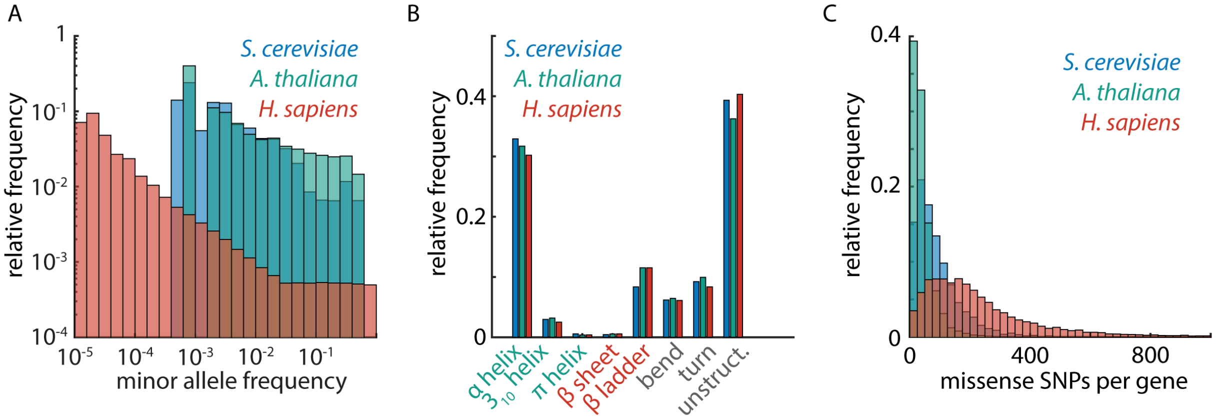


**Extended Data Figure S4. To accompany Figure 4.** (A) Relative frequency of minor allele frequencies in databases of natural genetic variation in *S. cerevisiae*, *A. thaliana*, and *H. sapiens*, as indicated. (B) Relative frequency of protein secondary structural elements in *S. cerevisiae*, *A. thaliana*, and *H. sapiens*, as indicated. (C) Relative frequency of total number of observed missense variants per gene in databases of natural genetic variation in *S. cerevisiae*, *A. thaliana*, and *H. sapiens*, as indicated.

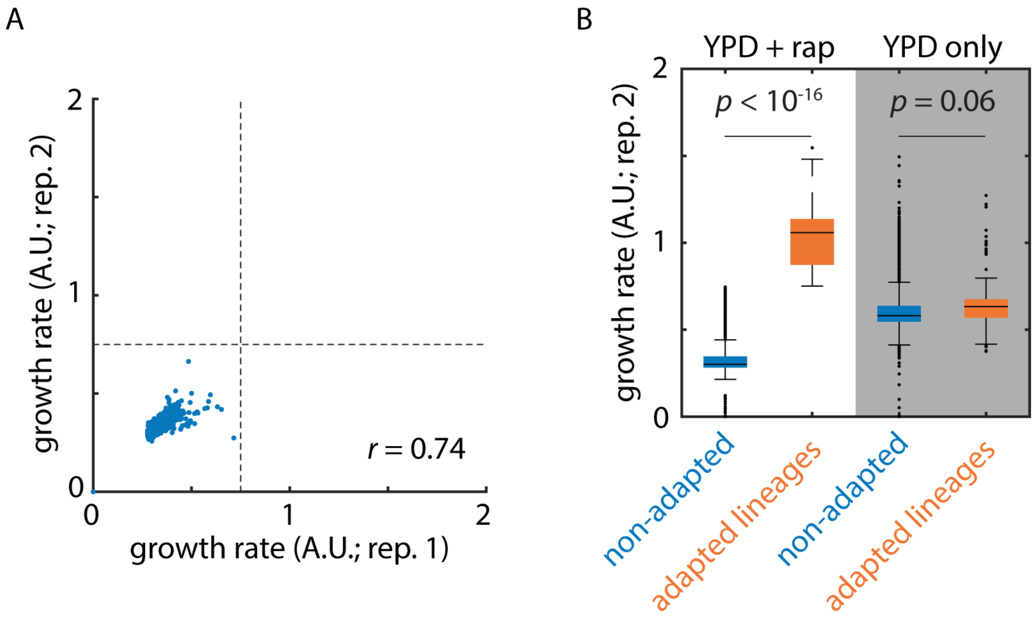


**Extended Data Figure S5. To accompany Figure 5.** (A) Replicability of estimated growth rates in rapamycin after archiving and revival of control lineages passages in the absence of rapamycin. (B) Estimated growth rates in rapamycin (left) and in the absence of drug (right) for adapted (blue) and non-adapted (orange) lineages. *p* values by *t* test.

| **Isolate** | **Chr** | **Pos** | **Mutation** | **Allele frequency** |
|---|---|---|---|---|
| 1 | XIV | 372039 | $\text{Fpr1}^{\text{I63T}}$ | 98.7% |
| 2 | XIV | 371998 | $\text{Fpr1}^{\text{E77K}}$ | 99.3% |
| 3 | XIV | 372144 | $\text{Fpr1}^{\text{L28*}}$ | 98.4% |
| 4 | | | no fixed mutations | |
| 5 | | | no fixed mutations | |
| 6 | | | no fixed mutations | |
| 7 | VII | 471457 | $\text{Pdr1}^{\text{I281N}}$ | 99.3% |
| 8 | XIV | 372067 | $\text{Fpr1}^{\text{Q54*}}$ | 100% |

**Table S1.** SNPs identified in rapamycin-adapted isolates selected for whole-genome sequencing.

| | | | | | | |
|---|---|---|---|---|---|---|
| M1I (x2) | E6* | **L28*** | **Q54* (x3)** | Q61* (x2) | W66* (x4) | L73* (x3) |
| E77* | E109* (x2) | | | | | |

**Table S2.** Presumed loss-of-function mutants identified in pooled, targeted sequencing of the *FPR1* locus. Mutations also identified in whole-genome sequencing [**Table 1**] in **bold**. Mutations isolated in the 1991 identification of rapamycin-resistant *FPR1* mutants[34] underlined.
